# Dynamic networks of fighting and mating in a wild cricket population

**DOI:** 10.1101/475087

**Authors:** David N. Fisher, Rolando Rodríguez-Muñoz, Tom Tregenza

## Abstract

Reproductive success is often highly skewed in animal populations. Yet the processes leading to this are not always clear. Similarly, connections in animal social networks are often non-randomly distributed, with some individuals with many connections and others with few, yet whether there are simple explanations for this pattern has not been determined. Numerous social interactions involve dyads embedded within a wider network. As a result, it may be possible to model which individuals accumulate social interaction through a more general understanding of the social network’s structure, and how this structure changes over time. We analysed fighting and mating interactions across the breeding season in a population of wild field crickets under surveillance from a network of video cameras. We fitted stochastic actor-oriented models to determine the dynamic process by which networks of cricket fighting and mating interactions form, and how they co-influence each other. We found crickets tended to fight those in close spatial proximity to them, and those possessing a mutual connection in the fighting network, and heavier crickets fought more often. We also found that crickets who mate with many others tended to fight less in the following time period. This demonstrates that a mixture of spatial constraints, characteristics of individuals and characteristics of the immediate social environment are key for determining social interactions. The mating interaction network required very few parameters to understand its growth and so structure; only homophily by mating success was required to simulate the skew of mating interactions seen in this population. This demonstrates that relatively simple, but dynamic processes can give highly skewed distributions of mating success.

## Introduction

Organisms engage in social interactions when they mate, fight, cooperate and compete for resources with conspecifics (Frank 2007). Interactions such as these influence an individual’s fitness and allow it to influence the fitness of others (Formica et al. 2012; Royle et al. 2012; Wey et al. 2013). Social interactions can therefore play a key role in ecological and evolutionary processes. Furthermore, these interactions are temporally dynamic, as individuals change interactions partners over time (Blonder and Dornhaus 2011; Blonder et al. 2012). This may influence the rate at which individuals encounter potential mates or competitors, the rate of opportunities for pathogen and information transmission, and the opportunities for different social strategies (Pinter-Wollman et al. 2013). How individuals accumulate social interactions is therefore key for several aspects of their fitness.

Reproductive skew in wild populations is typically substantial, with many individuals achieving no or little success, while some individuals are highly successful (Keller & Reeve 1994; Clutton-Brock *et al.* 1997; Engh 2002; Frentiu & Chenoweth 2008; Ryder *et al.* 2009; Rodríguez-Muñoz *et al.* 2010; Thompson *et al.* 2011). This indicates a skew in social interactions, with some individuals having many mating connections, while most having very few or none. In fact such a skew is common across all sorts of social networks, where most individuals have few connections, while a small number of others are very well connected (Croft et al. 2008; Krause et al. 2014). Since both a network of social interactions and a set of mating interactions in a population arise from many dyadic interactions accumulating over time, this raises the possibility that similar processes give strong skews in mating success and social network connections. Not mutually exclusively, it is also possible that the accumulation of interactions in one context influences the interactions in the other context, so for example a high number of interactions in a grooming network leads to many connections in a mating network.

Networks with properties similar to real-world networks can be simulated by dynamic network growth models with few rules (Newman 2002; Ilany and Akçay 2016), indicating that a network’s structure can be directly depend on the dynamic processes that form it. Similarly, simple rules that individuals follow in relation to the movement of fellow group members can result in the apparently complex patterns displayed in murmurations of starlings or synchronised swimming in shoals of fish (Sumpter 2006; Rosenthal et al. 2015). Understanding individual-level decisions about interactions with other population members may therefore allow us to explain the structure and properties of whole groups, including the spread of mating interactions within a population.

To investigate these topics, we use dynamic social network analysis to explore how fighting interactions accumulate over time within the lifetimes of individually marked wild adult field crickets (*Gryllus campestris*) to give highly non-random social networks. We then looked at how networks of mating interactions co-change over time alongside the fighting networks, and how these two networks dynamically influence each other. We therefore assessed what processes underpinned the formation of these two networks, and so what could explain the skew in connections apparent in each (Figs. 1 & 2), as well as how they influence each other.

**Figure 1.**
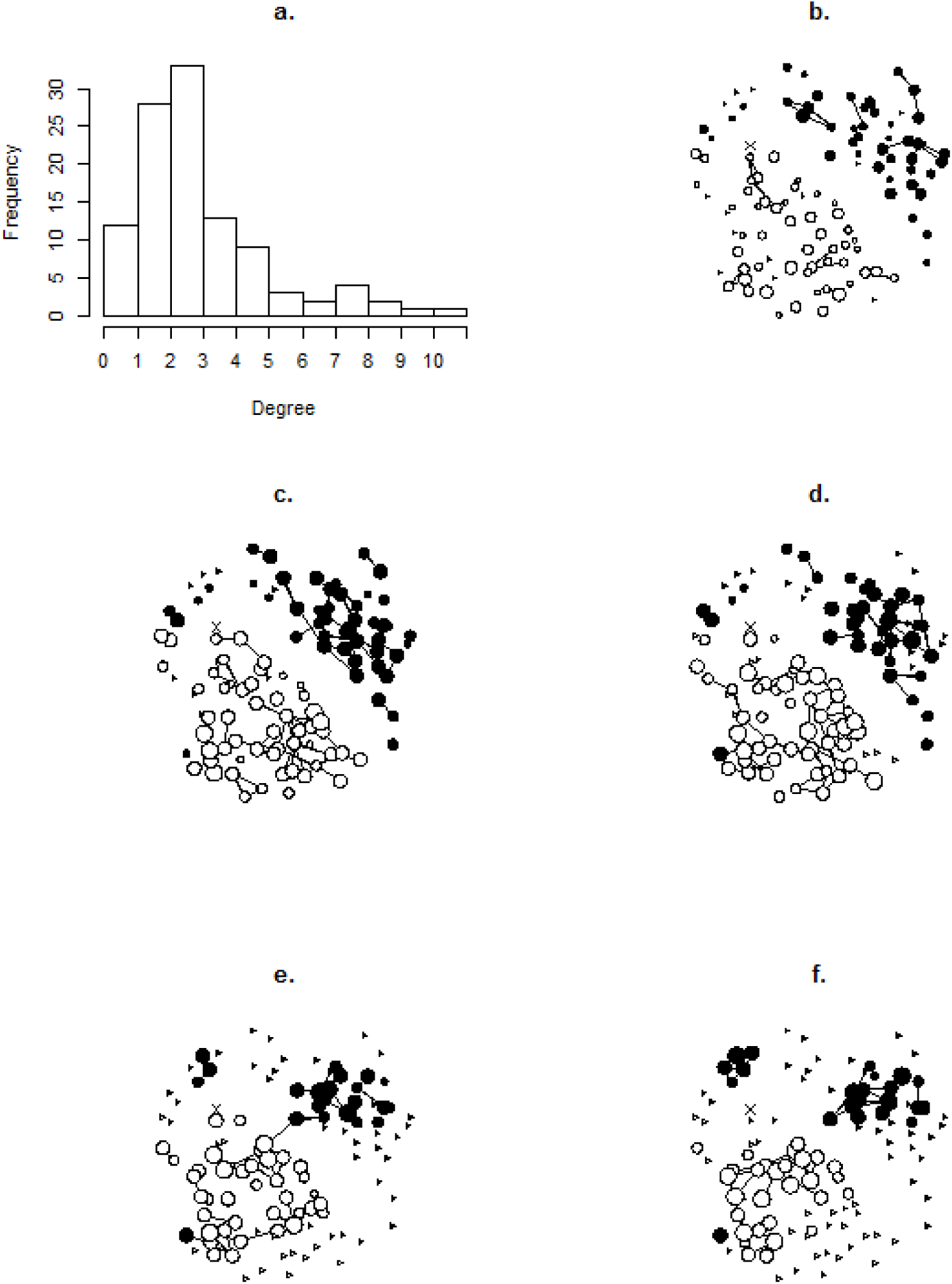
The degree distribution of the fighting network (a.), and a network plot for each of the five time-periods (b-f.). For the degree distribution all five time periods are aggregated to give the frequencies of the total number of different crickets an individual fought in 40 days. For the network plots, males are filled circles, and females open circles. The size of the circular nodes indicates the activity level of the individual (from 1-4) with individuals who were not alive during the time-period plotted as a small triangle. The position of an individual is the same in each plot, using a Fruchterman-Reingold algorithm (Fruchterman and Reingold 1991) based on an aggregation of all five time-periods. For illustrative purposes, the “X” indicates a female who fought two other individuals and recorded 3 leaves events (and so an activity level of 2) in the 1^st^ time-period, 1 fight and 4 leaves (activity = 2) in the 2^nd^ time-period, 0 fights and 9 leaves (activity = 3) in the 3^rd^ time-period, 0 fights and 12 leaves (activity = 3) in the 4^th^ time-period, and was dead for the 5^th^ time-period. Networks plotted using the R package “network” (Butts 2008).

## Methods

### Study site

The study site is located in a meadow in Northern Spain, see www.wildcrickets.org and Rodríguez-Muñoz *et al.* (2010; 2018) for further information. We used data collected in 2013 for this analysis. In the early spring we located each burrow and marked it with a unique identifier. In late April, just before adults start to emerge, we set out 124 cameras at random at burrows with an active juvenile cricket (nymph). This allowed us to record the exact moment of emergence for those adults, and all subsequent behaviour at the burrows. We directly monitored burrows that were without cameras daily or every other day and recorded the life stage and identity of the individual using the burrow. As nymphs only very rarely move among burrows, when there was an untagged adult at a burrow where on the previous days there had been a nymph, we could infer the emergence date for that adult. This allowed us to record accurate emergence dates for the vast majority of the population. Adults mate with members of the opposite sex, fight individuals of typically the same sex and hide from predators at these burrows, so by monitoring the burrows directly we capture the vast majority of relevant cricket behaviour. If we did not observe a cricket’s death, we estimated it as the day after it was last observed. A few days (mean ± standard deviation = 3.76 ± 2.81) after a cricket emerged as an adult, we trapped it (using a custom-built trap, see www.wordpress.com/crickettrapping for more details), and transported it to a laboratory adjacent to the field site. Here we weighed it and fixed a water-proof vinyl tag to its pronotum using cyanoacrylate glue. This allows the identification of individuals, and as far as we are aware does not affect their natural behaviour. After tagging the crickets, we released them back to the burrow they were trapped from, which we kept blocked in the meantime to prevent other animals, including other crickets, from usurping the burrow. We moved cameras away from burrows that hosted no cricket activity for two days to nearby ones where cricket activity had been directly observed or which showed signs of activity. As the season progresses there become more cameras than live adult crickets. This gives us very good information on behaviours over individuals’ entire adult lifetimes. In the centre of the meadow there is a weather station that takes measurements of rain fall and intensity of solar radiation every ten minutes (Vantage Pro 2, Davis instruments, California).

### Study organism

*G. campestris* is univoltine and adults are active in the months April – July following overwintering as nymphs in burrows they dig themselves. Once sexually mature, males start calling to attract mates, and both sexes move among burrows to search for mating partners. When encountering a member of the same sex at a burrow they will typically fight, with the loser leaving the burrow (Alexander 1961). While many male *and* female *G. campestris* achieve very low fitness, small males that sing frequently, and larger, long lived, and more promiscuous individuals of both sexes achieve higher lifetime reproductive success (Rodríguez-Muñoz et al. 2010). In *G. campestris,* reproductive success is strongly influenced by mating success (Rodríguez-Muñoz et al. 2010), although post-copulatory processes may have some influence (Bretman and Tregenza 2005; Bretman et al. 2009, 2011). Hence the use of accumulation of mating interactions as a proxy for the accumulation of fitness is reasonable.

### Modelling dynamic networks with SAOMs

We used stochastic actor-orientated models (SAOMs) to model the dynamic networks of mating and fighting interactions in our population of field crickets and therefore 1) determine processes that lead to the skew in connections in each and 2) determine how the networks influence each other. SAOMs allow the modelling of the change in individuals’ social interactions and behaviours over time, as influenced by individual or dyadic (depending on some aspect of the existing relationship between two individuals) effects and properties of the existing network structure (Steglich et al. 2006; Burk et al. 2007; Snijders et al. 2010). SAOMs can also be used to study transmission dynamics (e.g. Pasquaretta et al. 2016; Silk et al. 2017), and the effect of environmental factors on social interactions (e.g. Ilany et al. 2015). These models are therefore useful for testing a range of hypotheses of interaction in behavioural and evolutionary ecology (Fisher et al. 2017). We implemented our SAOMs in the R package “RSiena” (Ripley et al. 2015).

### Network construction

Initially we were interested in the fighting behaviour of individual crickets. We judged two crickets to have fought if there was any kind of aggressive interaction between them, which can be unidirectional. These fights typically occur immediately after a cricket arrives at a burrow at which there is already a member of the same sex. The loser will then leave the burrow. These fights are assumed to be over potential mating partners (Alexander 1961) or to provide access to the safety of a burrow. We split the season into nine eight-day time-periods, which gives a manageable number of time steps but also allows enough time for interactions to occur to prevent each time-period having a low frequency of interactions. To avoid exceptionally sparse networks we removed crickets (n = 58) who only fought a single other individual in a single time-period, leaving networks of 108 individuals. For each time-period we created a network, linking individuals if they fought at least once in that time-period (hence the networks were binary). If an individual was not alive during a time-period we entered “structural zeroes” for all its potential interactions. These indicate that interactions with that individual could not have taken place, preventing the lack of interaction from informing parameter estimates (Ripley et al. 2015).

For the mating network, we linked crickets in a network if they mated at least once in the eight-day period, similar to the fighting network (again a binary network). We added structural zeroes for all potential interactions between individuals of the same sex, as such interactions in that network were impossible. This was input into a SAOM alongside the networks of fighting behaviour, as we expected them to influence each other. We limited both networks to crickets who mated or fought more than one other cricket or mated or fought in more than one time-period, giving networks of 113 crickets, a slightly larger subset of the population than used previously, again to prevent exceptionally sparse networks. For both networks, if an individual was not alive during a time-period we entered structural zeroes for all its potential interactions.

### Network analysis

Unless otherwise stated, we used the same method and rationale as outlined in Fisher *et al*. (2017) and that article’s supplementary materials. We initially had nine eight-day time-periods. However, in the first two and last two time-periods there were not enough social interactions to investigate the processes that influence their change, so we did not use them, leaving the middle five time-periods (spanning 20/5/2013-28/6/2013). Terms are considered significant at the 95% if the absolute value of “estimate / standard error” was > 2 (Burk et al. 2007; Ripley et al. 2015). Bellow we explain the modelling process for each of the networks.

For the fighting network we used a forcing model (model type 2), where one individual dictates whether a tie is created or dissolved (Ripley et al. 2015), as for fighting a cricket can simply attack another or leave the area when they both meet. The initial SAOM for fighting behaviour contained rate parameters for each time-period and the effects of “density” (the tendency for individuals to be connected to all others in the network, typically negative as networks are generally sparse) and “triadic closure” (the tendency for individuals to form connections with those they share a mutual connection with, typically positive as individuals interact with those they share a mutual connection with). We tested this for satisfactory goodness-of-fit (GOF) with three network statistics: degree distribution (the frequencies of the different numbers of unique connections possessed by crickets in the networks), geodesic distribution (the frequencies of the different shortest path lengths in the networks) and the triad census (the frequencies of each set of three crickets that possessed 0, 1, 2 or 3 links among them, c.f. Ilany et al. 2015). These are chosen as they are commonly calculated network statistics, but their values are not defined by any of the parameters in the model (Ripley et al. 2015). The observed network statistics were not different from the network statistics of the set of networks generated by the model fitting process (p = 0.281, 0.399 & 0.994 for the GOF tests for degree distribution, geodesic distribution and the triad census respectively), indicating a satisfactory fit had been achieved. We therefore began adding terms of interest. After adding a term, we ran the model until it achieved convergence, and assessed the GOF. If the GOF had worsened we removed the newly-added term(s) before continuing, otherwise it/they were retained.

First, we added the individual covariate of sex, and two parameters, one for sex affecting the number of interactions an individual has, and one for interactions occurring depending on the sex of both individuals. The former term models the tendency for members of one sex to fight more often than members of the other sex, which we expect to have little effect based on previous results (Fisher et al. 2016a,b). The latter term models the tendency for crickets to predominantly fight members of the same sex as themselves, which we expected to be a strong effect as fights between males and females are exceptionally rare. We next added a changing dyadic covariate of distance, which was the Euclidean distance between each pair of crickets at the start of the time-period. This models the extent to which crickets nearer each other are more likely to interact than those further away. As a SAOM models the transitions between networks, rather than the structure of the networks themselves, we entered four instead of five measures of distance for the four transitions. We then added the constant covariate of individual mass (g), and its effect on the number of connections and individual acquired, and the interaction between the mass of each individual and its potential associates. We expected heavier crickets to fight more often (Dixon and Cade 1986), and crickets to avoid fighting much heavier individuals (Arnott and Elwood 2009). We next added two effects for weather: the total amount of rainfall (cm) and the intensity of solar radiation (Watts/m^2^) in each time-period. These were predicted to increase and decrease the frequency of social interactions respectively, as they have concurrent effects on movement around burrows (Fisher et al. 2015). Each individual is scored as being exposed to the same amount of rainfall and solar radiation in each time-period. Each term did not worsen the GOF of the model (not shown) and so were retained. This is the final model for the fighting network dynamics.

### Mating and fighting networks

To simultaneously analyse mating and fighting we entered the five mating networks alongside the five fighting networks into a SAOM. We used a unilateral initiative and reciprocal confirmation model (model type 3; Ripley *et al.* 2015), since for mating, both crickets need to be receptive for it to occur. This model initially includes the effects of density and triadic closure for both networks. We removed the effect of triadic closure from the mating network, as it is impossible in this network (as the third interaction in the triad would have to be a same-sex mating). Once this model converged, we began adding terms. The GOF for the mating network was not initially satisfactory (p = 0.019, 0.041 & 0.008 for the GOF tests for degree distribution, geodesic distribution and the triad census respectively) so we added the effect of “degree assortativity” for the mating network. If significant and positive, this effect indicates that individuals with many associations preferentially interact with other individuals with many associations. This possibly represents mutual mate choice, something we have found inferential evidence for previously (Fisher et al. 2016a). This model converged, and achieved satisfactory GOF (p = 0.413, 0.612 & 1.00 for the GOF tests for degree distribution, geodesic distribution and the triad census respectively), so we began adding terms of interest.

We first added the changing dyadic covariate of distance for both networks, calculated in the same way as previously of the fighting network. We next added the effect of mass for both networks, and the interaction between the mass of two potential associates for the mating network. The latter effect was not added for the fighting network in this two-network model, as the prior results indicated it was not important (Table 1), and we wished to avoid over-parameterising the model. We expected mass to be positively related to mating interactions, but for the interaction to not be important, as individuals of all sizes may prefer larger, presumably more fecund individuals (e.g. Aquiloni & Gherardi 2008; Baldauf *et al.* 2009). We also added the effects of rainfall and solar radiation for the mating network. These were not added for the fighting network as previous results indicated they were not important (Table 1).

**Table 1.** Results for the SAOM for the fighting network. Shown are the effect estimates, standard errors, convergence scores and the t-statistics (estimate / standard error). Effects are considered significant at the 95% level when the absolute t-statistic is greater than two. Such effects (aside from the rate parameters) are highlighted in bold. Rate parameters in a SAOM with only one dependent network are calculated rather than estimated, so convergence scores are not given here.

| Effect name | Estimate | Standard error | Convergence | t-statistic |
| --- | --- | --- | --- | --- |
| *Rate of change (period 1)* | 3.300 | 1.130 | NA | 2.921 |
| *Rate of change (period 2)* | 2.169 | 0.373 | NA | 5.811 |
| *Rate of change (period 3)* | 1.040 | 0.191 | NA | 5.456 |
| *Rate of change (period 4)* | 1.913 | 0.456 | NA | 4.200 |
| ***Density*** | **-4.519** | **0.355** | **0.057** | **-12.739** |
| ***Triadic closure*** | **0.861** | **0.217** | **0.024** | **3.959** |
| ***Distance*** | **-0.159** | **0.018** | **-0.075** | **-8.790** |
| ***Sex*** | **-0.414** | **0.183** | **-0.031** | **-2.262** |
| ***Sex ego x Sex alter*** | **6.398** | **1.144** | **0.059** | **5.595** |
| ***Mass*** | **1.991** | **0.892** | **-0.003** | **2.232** |
| *Mass ego x mass alter* | -5.214 | 5.466 | -0.005 | -0.954 |
| *Rainfall* | 0.007 | 0.013 | 0.058 | 0.592 |
| *Solar radiation* | < 0.001 | < 0.001 | -0.025 | 1.500 |

**Table 2.** Results for the mating and fighting network SAOM used for the third (and to a lesser extent the first) question. Effects are considered significant at the 95% level when the absolute t-statistic is greater than two. Such effects (aside from the rate parameters) are highlighted in bold. The four rate-of-change parameters for the fighting network were fixed rather than freely estimated, hence their statistics other than the estimate are not provided (see Table 1).

| *Fighting network effects* | *Estimate* | *Standard*<br>*error* | *Convergence* | *t-statistic* |
| --- | --- | --- | --- | --- |
| *Rate of change (period 1)* | 3.300 | NA | NA | NA |
| *Rate of change (period 2)* | 2.169 | NA | NA | NA |
| *Rate of change (period 3)* | 1.040 | NA | NA | NA |
| *Rate of change (period 4)* | 1.913 | NA | NA | NA |
| ***Density*** | **-2.004** | **0.170** | **-0.067** | **-11.795** |
| ***Triadic closure*** | **0.862** | **0.221** | **-0.026** | **3.907** |
| *Distance* | -0.005 | 0.016 | 0.072 | -0.313 |
| *Sex* | -0.129 | 0.135 | 0.030 | -0.955 |
| ***Sex ego x Sex alter*** | **3.270** | **0.577** | **-0.076** | **5.672** |
| *Mass* | 0.997 | 0.714 | 0.056 | 1.396 |
| ***Popularity in mating***<br>***network*** | **-0.637** | **0.291** | **0.027** | **-2.185** |

| *Mating network effects* | *Estimate* | *Standard*<br>*error* | *Convergence* | *t-statistic* |
| --- | --- | --- | --- | --- |
| *Rate of change (period 1)* | 5.306 | 1.490 | 0.015 | 3.558 |
| *Rate of change (period 2)* | 3.829 | 1.018 | -0.009 | 3.761 |
| *Rate of change (period 3)* | 3.280 | 0.894 | 0.013 | 3.669 |
| *Rate of change (period 4)* | 3.664 | 1.657 | 0.007 | 2.212 |
| ***Density*** | **-1.605** | **0.118** | **-0.002** | **-13.609** |
| ***Degree assortativity*** | **0.158** | **0.066** | **-0.004** | **2.411** |
| *Distance* | -0.019 | 0.017 | 0.004 | -1.139 |
| *Mass* | -0.610 | 0.715 | -0.033 | -0.853 |
| *Mass ego x Mass alter* | -1.704 | 4.520 | -0.019 | -0.377 |
| *Rainfall* | 0.011 | 0.007 | -0.028 | 1.454 |
| *Solar radiation* | < 0.001 | < 0.001 | 0.001 | 0.343 |
| *Popularity in fighting network* | -0.026 | 0.185 | -0.033 | -0.138 |
| *Mutual partner* | 1.143 | 0.838 | -0.009 | 1.364 |
| ***Maximum Convergence Ratio = 0.146*** |  |  |  |  |

We then added terms relating to the co-evolution between the networks. The first of these was the effect of “across-network popularity”, where the number of an individual’s connections in one network influences its number of connections in the other network. We actually added two effects here, one for the mating-networks’ effect on the fighting networks, and then the effect in the opposite direction. We expect this to be positive, as individuals engaging in many fights are assumed to be doing so to gain access to many mating partners, while individuals mating with many partners presumably are also encountering many rivals to fight with. We finally added the “mutual partner” effect, from the fighting network to the mating network. This models the possibility that two individuals that fight are then more likely to share a mutual connection in the mating network. We have previously found that two males who fight are also typically in sperm competition (Fisher et al. 2016a) so we expect this effect to be positive. This was our final model.

## Results

### Fighting network

From the final model of fighting we found significant effects of density, triadic closure, the spatial distance between two individuals, an individual’s mass and both the main effect of sex and the interaction between the sexes of two potential associates (Table 1). The density effect was strongly negative, indicating that crickets tend not to be connected to all other crickets, and so the network is relatively sparse, like most social networks (Snijders et al. 2010). Triadic closure was positive, indicating that the presence of a mutual connection increased the chances of two crickets fighting. This was true even when accounting for the effect of distance between individuals, which negatively influenced their tendency to have interactions. The sex effect was negative, indicating that males fought fewer other individuals than females, while the interaction between the sex of one cricket and the sex of another was strongly positive, indicating fights are predominantly intra-sex. Heavier crickets fought more other crickets, again as predicted, but individuals did not avoid fighting those of greatly different weight to them (the interaction between the mass of an individual and the mass of its potential fighting partner was not important). The weather variables did not influence the fighting network.

### Mating and fighting networks

In the SAOM for the mating and fighting networks, all the significant effects from the previous analysis of the fighting network were in the same direction as before, although the effects of sex, distance and mass were not significant (Table 3). This possibly indicates a lack of power in this analysis. The effect of across network popularity from the mating network to the fighting network was significantly negative, indicating that individuals who mate with many others fight fewer other crickets in the next time-period.

For the mating network, the density effect was strongly negative as for the fighting network, again indicating the mating network is sparse much like other social networks. The effect of degree assortativity was positive, indicating that promiscuous males mated with promiscuous females. Otherwise no effects were significant, but since there is a lack of power in this analysis we will mention the following effects that were close to significance (|estimate / standard error| >1). Increasing distance decreased the likelihood of mating interactions, while rainfall increased their likelihood. The mutual partner effect was positive, suggesting that crickets who are connected in the fighting network tend to be more likely to share a mutual connection in the mating network. Neither the main effect of mass nor the interaction between mass of the male and female were important, nor was the effect of solar radiation and the effect of popularity in the fighting network.

**Figure 2.**
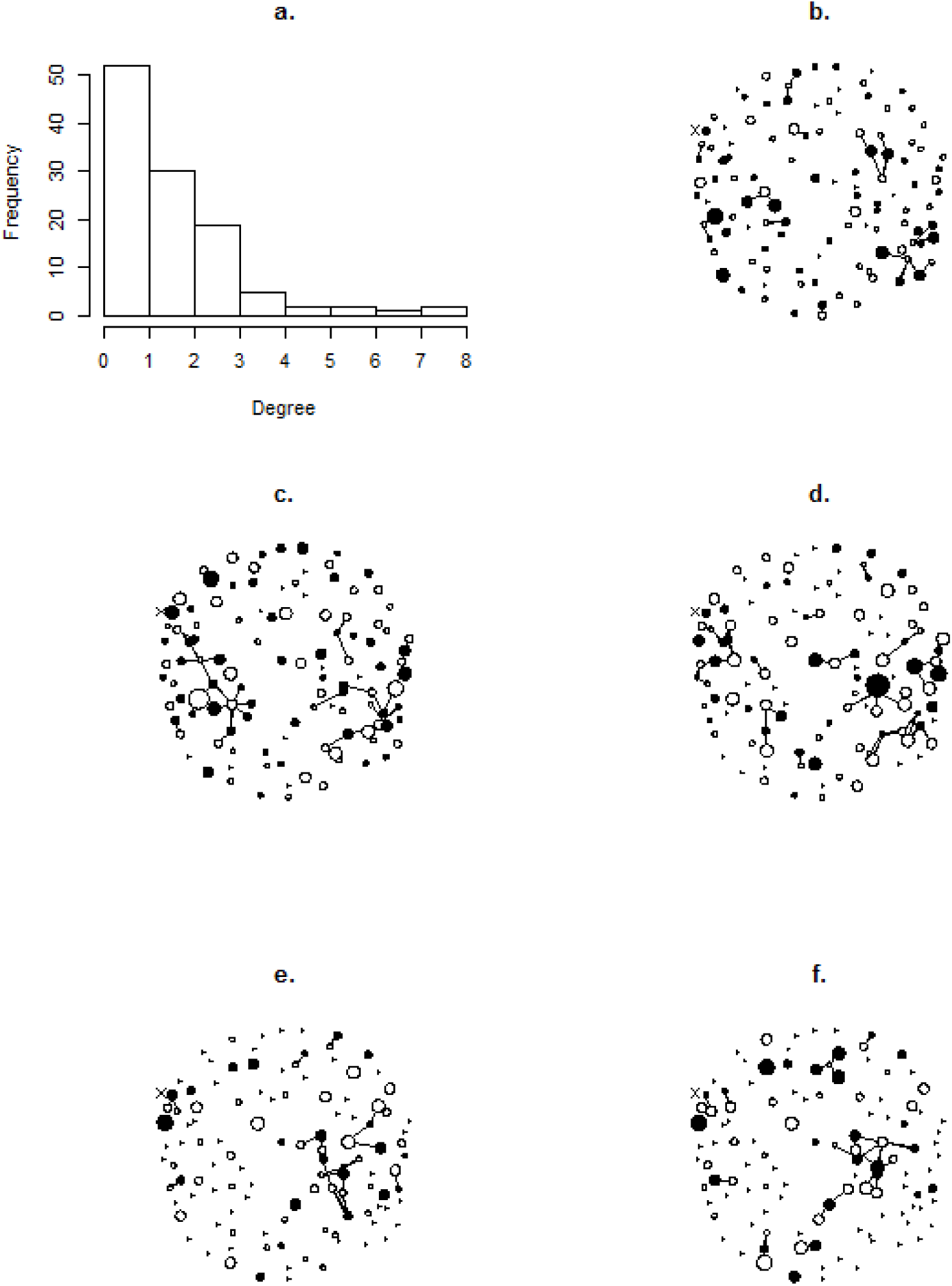
The degree distribution of the mating network (a.), and a network plot for each of the five time-periods (b-f.). For the degree distribution all five time points are aggregated to give the frequencies of the total number of different crickets an individual mated with over 40 days. For the network plots, males are filled circles, females open circles. The size of the circular nodes indicates the degree in the fighting network of that individual in that time-period. Individuals who were not alive in the time-period are plotted as small triangles. The position of an individual is the same in each plot, using a Fruchterman-Reingold algorithm (Fruchterman and Reingold 1991) based on an aggregation of all five time-periods. For illustrative purposes, the “X” indicates a male that had no matings and a single fight in the first time-period, 1 mating and 3 fights in the 2^nd^ time-period, no matings and 1 fight in the 3^rd^ time-period, 2 matings and 2 fights in the 4^th^ time-period, and 1 mating and no fights in the 5^th^ time-period. Networks plotted using the R package “network” (Butts 2008).

## Discussion

Overall, using two SAOMs we could recapture the skew in social interactions that occurs in both the fighting and mating networks. This demonstrates that individual mass, physical distances and the presence of mutual connections with males and females all influence the accumulation of fighting interactions in wild crickets, and lead to a skew in social interactions that is common to the vast majority of social networks. We were also able to recapture the skew in interactions in the mating network. This demonstrated that a relatively simple process, the assortment of individuals by their existing number of connections, can lead to the kind of skew in mating success that is very commonly observed in nature. Furthermore, we identified how the mating interaction network influences the fighting interaction network. This shows how social interactions in one context can influence the accumulation of interactions in another context. We now deal with each of our results in more detail.

### Interactions in the fighting network

We found that males fought fewer different individuals than females (see also: Fisher et al. 2016b). This does not necessarily mean that females are more aggressive; in this species, while both sexes engage in active mate searching (Hissmann 1990), typically it is females that move between burrows, while males sit and sing to attract them. Females are therefore more likely to encounter several different females as they are moving among burrows, and so be involved in an aggressive interaction with them. Males may engage in repeated fights with the same individuals, especially if they are calling from nearby burrows. Fighting amongst males does not decrease the intensity of sperm competition between them (Fisher et al. 2016a), and since fights have inevitable energetic costs and carry the risk of injury, male fights may not bring sufficient sexually selected benefits to drive combat with many different rivals.

The effect of spatial distance was negative, as expected. In many species individuals will associate more with those close to them, so controlling for spatial proximity when attempting to detect genuinely socially driven associations is important (Whitehead and James 2015). However, the relationship is likely to be bidirectional for many species, with space-use influencing who you interact with and animals moving based on the results or potential consequences of social interactions (Cantor et al. 2012; Webber and Vander Wal 2018). This makes simply “controlling” for space use problematic when space use itself may be an expression of social behaviour.

Heavier crickets fought more different individuals. This may suggest that fighting is a condition dependent strategy (Luttbeg and Sih 2010) or that heavier individuals employ a different social strategy that involves attempting to dominate their rivals (Hack 1997; Brown et al. 2006). The interaction between the mass of an individual and its potential associates was however not important. This may reflect how we only modelled the occurrence of fights, not who won. It may well be that crickets of different sizes will encounter each other at a burrow and interact aggressively, and then the size difference influences the outcome.

Finally, we found no link between the weather variables and frequency of fighting behaviour. We consider it unlikely that rain and solar radiation do not influence cricket social interactions, as crickets’ activity levels on a given day are influenced by the amount of rain and solar radiation (Fisher et al. 2015). Instead, we suspect that the eight-day periods we selected were too coarse a scale to detect these fine-scale behavioural responses. Ilany *et al.* (2015) found that wetter years lead to more sparse spotted hyena (*Crocuta crocuta*) social networks using a SAOM, so relationships between environmental and network characteristics can be detected with this approach in some systems.

### Interactions in the mating network

After adding the term of degree assortativity, we were successful in simulating the mating network, including a highly skewed pattern of mating success. Reproductive skew is ubiquitous in natural populations (Keller & Reeve 1994; Clutton-Brock *et al.* 1997; Engh 2002; Frentiu & Chenoweth 2008; Ryder *et al.* 2009; Rodríguez-Muñoz *et al.* 2010; Thompson *et al.* 2011) and helps provide the variation in fitness necessary for adaptation. It would be very interesting to know to what extent other mating systems can be modelled in this manner, and whether the processes of degree assortativity is as important in other mating systems as it is in the crickets.

Lifetime reproductive success is correlated with number of mating partners in this species (Rodríguez-Muñoz et al. 2010). Therefore, assortment by promiscuity may indicate mutual mate choice or assortment by “quality” (Aquiloni and Gherardi 2008; Baldauf et al. 2009), which could increase the variance in reproductive success in the population if high-fecundity individuals pair. However, as males with many mating partners mate with more promiscuous females, they face increased sperm competition for each ovum of females they mate with. This will reduce the variance in reproductive success among-males (Sih et al. 2009). Both the main effect of mass and the interaction between the mass of an individual and the mass of its potential mating partners was not related to links in the mating network, suggesting mating partner choice is not based on mass. Instead, chemical cues such as cuticular hydrocarbons are likely to be important in mediating partner choice between closely related species (Tyler et al. 2015), so may play a role here.

Only degree assortativity was needed to get a satisfactory GOF for the mating network, perhaps suggesting that the mating system is quite simple and beyond these few terms only stochasticity plays an additional role in determining its structure. This would be troubling given the amount of effort that is devoted to understanding patterns of mate choice and sexual selection in the wild. However, there is the potential for a lot of different behavioural processes to be contained within the effect of degree assortativity, such as the trait(s) crickets are using for mate choice and the processes that generate variation in these traits that cannot be exploited by “cheats” who do not signal honestly. Additionally, we have only modelled the choice of mating partners, not the frequency of mating with a particular partner in an eight-day period, as we used binary networks. Therefore, there is likely variation in preference among mating partners that we are ignoring, which could influence fitness as frequency of copulation is likely related to share of paternity (Parker 1970; Simmons 1987).

We found that spatial distance did not significantly influence the mating network. This surprising result could stem from a number of sources. A lack of power as suggested earlier may have prevented us from detecting a biologically important effect. Alternatively, this may reflect the fact that there are many crickets near each other that do not mate. In general, if the choice of mates for an individual in a population is not limited to its immediate neighbours, simple models for population-level processes such as partner choice or sexual disease transmission that do not explicitly account for spatial constraints may be more accurate than thought (Patterson et al. 2008). The weather variables were also not important, but we hesitate to make conclusions about this since it may stem from looking at too coarse a scale as suggested above for the fighting network.

Individuals with more mating partners tended to have fewer fighting partners at the next time step. This seems to contrast with previous results that the involvement in fighting and sperm competition is positively correlated (Fisher et al. 2016a). However, these results are compatible if we consider the dynamic nature of the new result. Crickets over their lifetimes may show positive correlations between involvement in different types of competition, perhaps due to links to “quality” or differences in lifespan, but at any given time they may not be able do both (perhaps due to energetic constraints), creating a negative relationship between adjacent time steps. Furthermore, crickets that shared a mutual connection in the mating network were more likely to fight. This seems a direct response to the threat of sperm competition, as we have found previously (Fisher et al. 2016a). Crickets have flexible mating systems where they are involved in pre- and post-copulatory competition (Buzatto et al. 2014), so they are adapted to both physical contests and sperm competition. In other systems, where males can monopolise access to females through physical domination, we would not expect to see such a pattern.

## Conclusions

We have analysed how networks of fighting and mating interactions between crickets accumulate over time, and therefore arrived at a holistic understanding of how these networks come to be structured. By demonstrating that various individual and network-based factors influence social interactions, we have helped link social network analysis to existing theory on dominance interactions and sexual selection theory. These factors, along with stochastic processes, produced networks with a skewed degree distribution that mirrors the observed skew in social interactions and reproductive success in the population, suggesting these a network approach is an appropriate way to model these systems. We hope this stimulates others to use approaches such as this to gain more complete understanding of complex animal social systems.

## Acknowledgements

We thank Paul Hopwood, Alex Thornton and Andrew Jackson for comments that improved this manuscript. We also thank Luke Meadows and Carlos Rodríguez del Valle for assistance with data collection.

## Author contributions

All authors conceived of the research questions, contributed to study design and collected data. RRM & TT initiated the long-term study. DNF drafted the manuscript and conducted the data analysis. All authors approved of the final manuscript for submission.

## Funding statement

Funding for this study was provided by the Natural Environment Research Council (NERC); studentship: NE/H02249X/1; standard grants: NE/E005403/1, NE/H02364X/1, NE/L003635/1, NE/R000328/1.

